# RNA Pol II antagonises mitotic chromatin folding and chromosome segregation by condensin

**DOI:** 10.1101/2023.08.08.552486

**Authors:** Jeremy Lebreton, Léonard Colin, Elodie Chatre, Pascal Bernard

**Affiliations:** ENS de Lyon, Univ Lyon, 46 allée d’Italie, F-69007, Lyon, France; CNRS Laboratory of Biology and Modelling of the Cell, UMR 5239, ENS de Lyon, 46 allée d’Italie, F-69007, Lyon, France; Lymic-Platim, Univ Lyon, Université Claude Bernard Lyon 1, ENS de Lyon, CNRS UAR3444, Inserm US8, SFR Biosciences, 50 Avenue Tony Garnier, F-69007 Lyon, France

**Keywords:** condensin, SMC complexes, loop-extrusion, mitotic chromosome assembly, transcription, transcription-termination

## Abstract

Condensin shapes mitotic chromosomes by folding chromatin into loops but whether it does so by DNA-loop extrusion remains speculative. While loop-extruding cohesin is stalled by transcription, no conclusive evidence has been provided regarding the impact of transcription on condensin despite its conserved enrichment at highly expressed genes. Using degrons of Rpb1 or the torpedo nuclease Dhp1^XRN2^, we depleted or displaced RNAP2 on chromatin in fission yeast metaphase cells. We show that RNAP2 does not load condensin on DNA but instead retains condensin and hinders its ability to fold mitotic chromatin and to support chromosome segregation, consistent with the stalling of a loop-extruder. Transcription termination by Dhp1 limits such a hindrance. Our results shed a new light on the integrated functioning of condensin and we argue that a tight control of transcription underlies mitotic chromosome assembly by loop-extruding condensin.

## Introduction

During mitosis, chromatin is reshaped into rod-shaped chromosomes in preparation for the accurate segregation of sister chromatids in anaphase. The assembly and maintenance of mitotic chromosomes is driven by the nonhistone protein complex condensin^1^, whose deficiency manifests in anaphase by the stereotypical formation of chromatin bridges^2–4^. Condensin has two variants, named condensin I and II, with the latter being lost several times during evolution^1^. Budding and fission yeasts only have a single complex similar to condensin I in term of primary amino acid sequence. Current data indicate that yeasts and vertebrates condensins shape mitotic chromosomes by folding chromatin into loops^5–8^, but the underlying mechanisms remain incompletely understood.

Condensins belong to the conserved family of SMC (Structural Maintenance of Chromosomes) genome organizers, which in eukaryotes includes cohesin and SMC5/6^1^. SMC protein complexes are composed at their core of two SMC-ATPases and a kleisin subunit that together form a ring, which associates with additional regulatory subunits^1^. A prevalent model proposes that SMCs drive intrachromosomal 3D contacts by processively extruding loops of DNA^9^. Condensins, cohesin and SMC5/6 have been observed extruding naked DNA into loops in an ATP-dependent manner in *vitro*^10–14^. Hi-C studies have further shown that, in addition to mediating sister-chromatid cohesion, cohesin organises chromatin into loops and intrachromosomal topologically associated domains (TADs) during interphase^15,9^, while condensin-mediated loops enlarge during mitosis^5,6,8^. Although loop-extrusion *per se* has not been directly observed *in vivo*, a large body of studies support the idea that loop-extruding cohesin is halted by DNA-bound proteins such as pairs of convergent CTCF or active RNA polymerases in the context of chromatin^9,16–19^. However, similar experimental evidence remains scarce for eukaryotic condensins^20,21^. An alternative non-mutually exclusive model, namely diffusion capture^22^, proposes that condensin shapes mitotic chromosomes by stabilizing random 3D contacts between its binding sites, either by capturing two DNA molecules inside its ring or through condensin-to-condensin contacts. Biophysical simulation recapitulates features of budding and fission yeast mitotic chromosomes^22,23^, and experimental evidence indicates that condensin-condensin interactions underlie chromosome formation by a loop-extrusion independent mechanism in Xenopus egg extracts^24^. Thus, the mechanism(s) by which condensins fold chromatin into loops during mitosis remain(s) unclear. To gain in functional understanding of condensin, we sought to compare its interplays with RNA polymerases with those of cohesin, taking advantage of the fact that transcription remains active during mitosis in fission yeast^25^. Chromatin immunoprecipitation coupled with high throughput sequencing (ChIP-seq) studies performed from yeasts to human have revealed a same broad basal association of condensins along the genome punctuated by peaks of high occupancy at centromeres, telomeres and rDNA repeats, as well as in the vicinity of highly expressed genes of any class along chromosome arms^25–28^. While cis-acting factors localising condensin at rDNA repeats^29,30^, centromeres^31,32,30^ or telomeres^33^ have been identified, the mechanisms underlying the enrichment of condensin at highly expressed genes and its functional consequences remain poorly understood. In fission yeast, condensin accumulates at the 3’ends of genes highly-transcribed during mitosis^25^. Mouse and human condensin II are also enriched at active genes in cycling cells^34^. Even chicken and human condensin I, which bind transcriptionally silent chromatin in mitosis, accumulate in the vicinity of promoters that were active in the previous G2 phase^25,27^. We reported that nucleosome eviction from gene promoters facilitates the loading of fission yeast condensin onto DNA in vivo^35^. It also has been shown in xenopus egg extracts, which lack transcription, that the general transcription factor TFIIH promotes the loading of condensin I and II by competing with nucleosomes^36^. The TATA binding protein Tbp1^TBP^, as well as sequence-specific DNA binding transcription factors, such as TFIIIC or Ace2, have been involved in the localisation of condensins at their target genes in yeasts and/or mammalian cells, but whether they act as a condensin loaders or positioning devices remains unclear^26,37,38,34^. The functional significance of condensin enrichment at highly expressed genes is further questioned by the finding that, in both chicken and fission yeast cells synchronized in mitosis, those sites exhibit only marginally increased frequencies of chromatin loops as compared to non-enriched loci^6,8^. It has been proposed that fission condensin and human condensin I bind to unwound DNA segments generated by transcription and reduce such structures to promote chromosome segregation during mitosis^25,39^. On the other hand, there is evidence that induction of transcription by either RNAP1 or RNAP2 of the 35S coding region of rDNA repeats antagonises condensin binding^40,41^ and we recently suggested that backtracked RNA polymerases constitute a barrier for condensin^21^. Thus, no clear picture emerges as to the functional significance of condensin enrichment in the vicinity of highly expressed genes.

Here, we used degrons alleles to rapidly deplete or displace RNAP2 on mitotic chromatin in fission yeast cells and assessed the consequences on condensin’s localization, chromatin folding in metaphase and accurate chromosome segregation in anaphase. In contrast to a previous study^25^, we found that RNAP2 transcription does not recruit fission yeast condensin, but instead retains it in cis. We further show that RNAP2 hinders both condensin-mediated chromatin folding in metaphase and condensin-dependent accurate chromosome segregation in anaphase. We argue that our results are best explained by the stalling of translocating condensin against a transcriptional barrier and provide indirect evidence for condensin shaping mitotic chromosomes by DNA-loop extrusion.

## Results

### Transcribing RNAP2 causes the accumulation of condensin at the 3’end of class II genes

Fission yeast condensin binds chromatin throughout mitosis and accumulates at mitotically transcribed genes, forming peaks of high occupancy notably at the 3’end of class II genes whose expression is maximal in the M-G1 phases^25^. To revisit the role played by RNAP2 in condensin’s localisation, we created an auxin-inducible degron of Rpb1 (Rpb1-sAID), the largest subunit of RNAP2. Fission yeast cells expressing both the kleisin subunit of condensin tagged with GFP (Cnd2-GFP) and Rpb1-sAID were arrested in metaphase, exposed to either auxin (OFF condition) or NaOH (solvent, ON condition) while maintaining the arrest, and chromosomal associations of both Cnd2-GFP and Rpb1-sAID were measured by calibrated-ChIP (cal-ChIP). This type of experimental approach was used for all metaphase arrests. Auxin induced a rapid (30 minutes) and acute depletion of Rpb1 while Cnd2-GFP level and mitotic indexes remained unchanged (Fig. S1A-B). Quantitative (q)PCR analysis of representative mitotically-expressed protein-coding genes confirmed their near-complete loss of Rpb1 upon auxin adjunction (Fig. S1C, upper panel, n = 3 biological replicates). Consistent with a previous report showing that a chemical inhibition of RNA polymerases reduces condensin occupancy^25^, Rpb1 depletion from those genes caused a strong reduction of Cnd2-GFP, while it remained unchanged at control, non-RNAP2, condensin binding sites (Fig. S1C, lower panel). These results confirm that RNAP2 impinges upon condensin localisation. However, Cnd2-GFP occupancy remained constantly above background after depletion of Rpb1 (Fig. S1C), suggesting that a fraction of condensin persisted on chromatin.

To assess the genome wide relevance of these results, cal-ChIP samples were pooled and processed for high throughput sequencing (cal-ChIP-seq). Ranking ChIP-seq signals at protein coding genes by mean normalized Rpb1 signal revealed that Cnd2-GFP association mirrored Rpb1 (Rpb1-ON, Fig. 1A). Auxin-mediated depletion of Rpb1 caused a strong reduction of Cnd2-GFP at the 3’end of most if not all protein-coding and non-coding genes transcribed by RNAP2 (Rpb1-OFF, Fig. 1A-B and Fig. S1D) while a subset of gene promoters retained Cnd2-GFP occupancy (Fig.1B and S1C lower panel). It has been reported that treatment with the RNA polymerase inhibitor 1,10-phenanthroline causes condensin to dissociate from chromatin^25^. In contrast, when we scored the full normalized amounts of Cnd2-GFP reads mapped to the genome, we observed no major difference between the Rpb1-ON and Rpb1-OFF conditions (Fig. 1C), suggesting that Cnd2-GFP did not dissociate from chromatin and might instead translocate away from Rpb1-depleted genes. Cal-ChIP-seq also validated that auxin adjunction reduced the binding of Cnd2-GFP neither at tDNAs (Fig. 1D), nor at rDNA repeats or the kinetochore assembly site of centromere 1 (Fig. S1E,F) where condensin localisation relies respectively on the transcription factor TFIIIC^26,42,34^ and monopolin^30^. The enrichment of fission yeast condensin at the 3’end of active class II genes is therefore dependent on Rpb1 in *cis*. Consistently, ∼70% of the condensin peaks identified in RPB1-OFF were associated with tDNAs, rDNA genes or long terminal repeats retrotransposons (Fig. S1G) for which specific factors position condensin^26,31,34,43^. The remaining 30% contained no feature previously associated with condensin.

**Figure 1.**
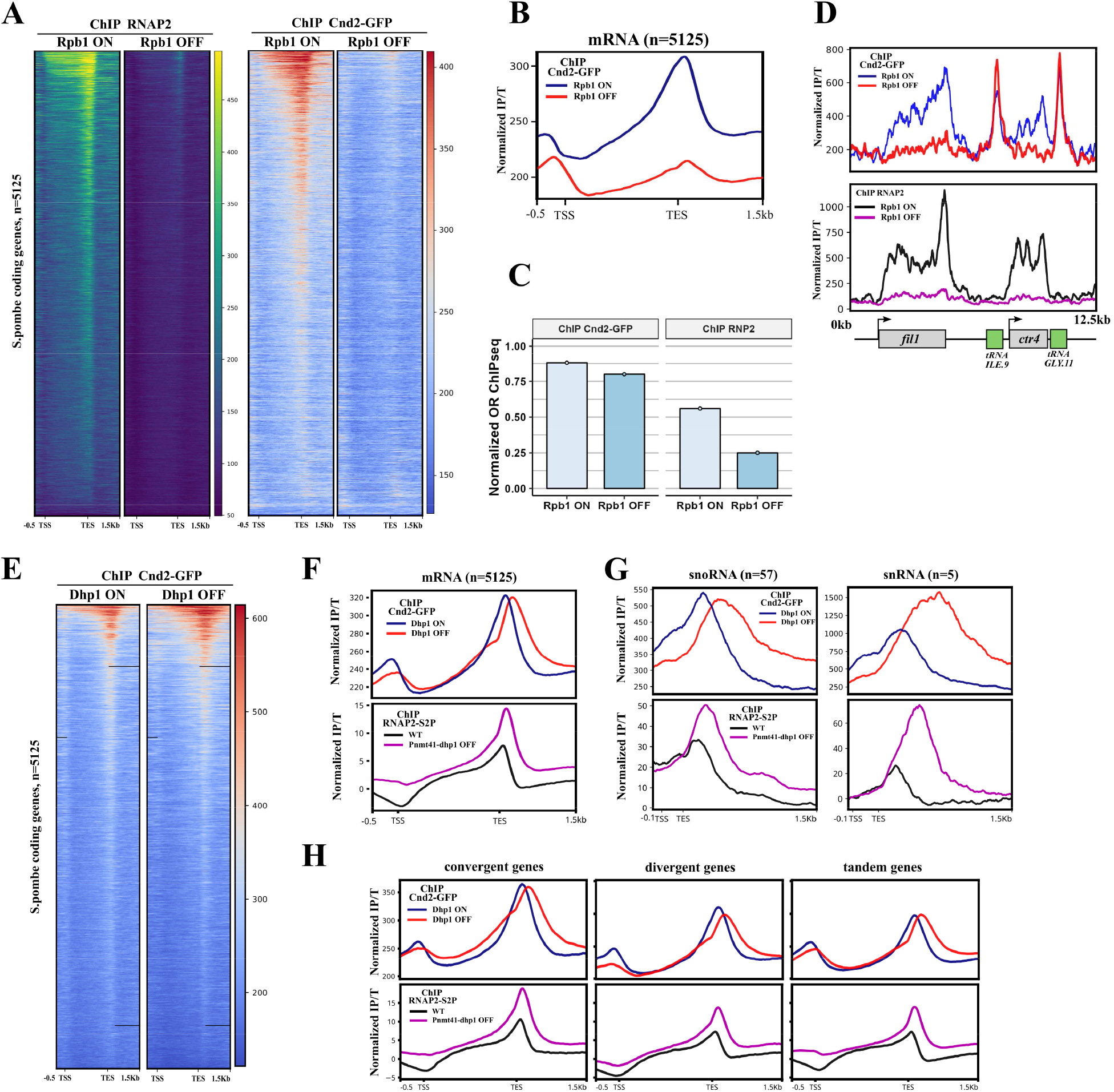
Active RNAP2 localises condensin in cis. (A) Heatmaps of normalized Rpb1 (8WG16) and Cnd2-GFP (A111-22) cal-ChIPseq signal at protein coding genes ranked by the mean Rpb1 signal, in NaOH (Rpb1-ON) or auxin-treated (Rpb1-OFF) *rpb1-sAID* metaphase arrested degron strains. The values represent the IP/T ratio of *S. pombe* calibrated by the *S. cerevisiae* SMC3-GFP IP (see Materials and Methods). (B) Mean metagene profile of the normalized Cnd2-GFP ChIPseq signal from (A). (C) Occupancy ratio (OR) of cal-ChIPseq from (A) in the indicated conditions. The OR is the ratio between the total amount of mapped reads in the IP and the Total of *S. pombe* normalized by the ratio between the amount of reads in the Total and the IP of *S. cerevisiae*. (D) Normalized Cnd2-GFP (top) and Rpb1 (bottom) cal-ChIPseq at chromosome III (810000- 822500) in Rpb1-ON and Rpb1-OFF. (E) Heatmap of normalized Cnd2-GFP cal-ChIPseq signal at protein coding genes ranked by mean strength, in NaOH (Dhp1-ON) or auxin-treated (Dhp1-OFF) *dhp1-sAID* metaphase arrested degron strains. The values represent the IP/T ratio of *S. pombe* calibrated by the *S. cerevisiae* SMC3-GFP IP (F-G) Metagene profile of the mean normalized cal-ChIPseq signal from (E) and the mean normalized RNAP2(S2P) cal-ChIPseq signal in WT or Pnmt41-dhp1-off from ref.44. (H) Metagene profile of the data in (F) grouped according to gene orientation.

If condensin is positioned by RNAP2 in mitosis, displacing RNAP2 on chromatin should shift condensin positions. To test this, we constructed an auxin inducible degron of Dhp1^XRN2^ (Dhp1-sAID), the torpedo ribonuclease involved in the termination of RNAP2 transcription^44^. Dhp1 loss of function in fission yeast causes active RNAP2 (S2P) to invade DNA sequences downstream of transcription end sites (TES)^44^. We depleted Dph1-sAID from metaphase arrests (Fig. S1A-B) and performed cal-ChIP-seq against Cnd2-GFP. Density heatmaps of read counts revealed an increase of Cnd2-GFP around the TES of a subset of protein-coding genes (Fig. 1E and S1H), but the most prominent phenotype was a shift of Cnd2-GFP peaks downstream the 3’end of genes (Fig. 1F). The shift was particularly obvious at sn(o)-RNA genes (Fig. 1G) and specific to class II genes (Fig. S1E-F). These results strengthen and extend a previous observation by ChIP-qPCR at two mitotically-transcribed protein-coding genes^39^. Larochelle et al. described a similar shift of Rpb1 cal-ChIP-seq signals in Dhp1-depleted fission yeast cells^44^. Reanalysing their data showed that the displacements of Cnd2-GFP upon Dph1- depletion mirrored those of Rpb1, both in orientation and in amplitude (Fig. 1F and 1G), arguing that condensin localisation responded to features associated with transcribing RNAP2. Since readthrough transcription can invade downstream genes, we analysed condensin localisation in Dhp1-ON vs OFF cells as a function of gene orientation (Fig. 1H). The shift of Cnd2-GFP signals upon Dhp1 depletion proved independent of gene orientation. However, convergent genes exhibited the highest condensin occupancy in Dhp1-ON and a specific increase of Cnd2-GFP occupancy in their body in Dhp1-OFF (Fig. 1H). This is consistent with our previous work suggesting that condensin is pushed by polymerases^21^ as converging reading-through RNAP2 may trap condensin. Thus, transcribing RNAP2 plays a role in *cis* in the accumulation of condensin at the 3’end of active class II genes.

### RNAP2 transcription does not recruit condensin onto DNA

Our finding that the steady state level of Cnd2-GFP on chromatin remained largely unchanged upon depletion of Rpb1 (Fig. 1C) appeared inconsistent with a preferential binding of condensin to unwound DNA generated by transcription^25,39^. To further investigate this, we took an orthogonal approach and performed photoactivated localisation microscopy and single particle tracking (SPT-PALM) of condensin in living fission yeast cells. A recent study has shown that SPT-PALM provides accurate measurements of the chromatin association of the related SMC5/6 complex in asynchronous fission yeast cells^45^. Using Cnd2 tagged with mEOS3.2 (Cnd2-mEOS3.2)^45^, we first characterized condensin behaviour in an otherwise wild-type (WT) background. Cells were arrested in metaphase, subsets of Cnd2-mEOS3.2 were photoconverted, imaged at a framerate of 33.6 ms, individual tracks were reconstituted and analysed with sptPALM viewer^46^. To identify metaphase cells, we used Cdc11-GFP to fluorescently-label spindle pole bodies (SPBs, the counterpart of centrosomes) and selected cells showing two SPBs separated by 2-3 µm (Fig. 2A). In WT, we observed two main populations of nuclear condensin characterized by their slow or fast molecular movements (Fig. 2B-C). The Cnd2-mEOS3.2 fast fraction showed a diffusion coefficient of around 0.5 μm^2^.s^−1^, reminiscent of the 0.7 μm^2^.s^−1^ of nucleosoluble Mcm4 in *S. pombe*^47^. By depleting the Cut14^SMC2^ subunit of condensin to detach Cnd2 from metaphase chromatin (Fig. 2D and S2A), we attributed the slow and fast Cnd2-mEOS3.2 signals to the chromatin-bound and nucleosoluble fractions of condensin, respectively (Fig. 2E). Next, we depleted Rpb1 from metaphase-arrested cells (Fig. S2A) and recorded Cnd2-mEOS3.2 movements. The fraction of slow, chromatin-bound, molecules of Cnd2-mEOS3.2 remained unchanged (Fig. 2F and S2B). In contrast, degrading Dhp1 in metaphase (Fig. S2A) slightly increased the steady state level of slow Cnd2-mEOS3 molecules (Fig. 2F and S2C) (see Discussion). Data analyses performed with another software (Spot-ON ^48^) provided identical results (Fig. S2D-E). These data confirm that RNAP2 transcription plays no role in the steady-state binding level of condensin to DNA *in vivo*.

**Figure 2.**
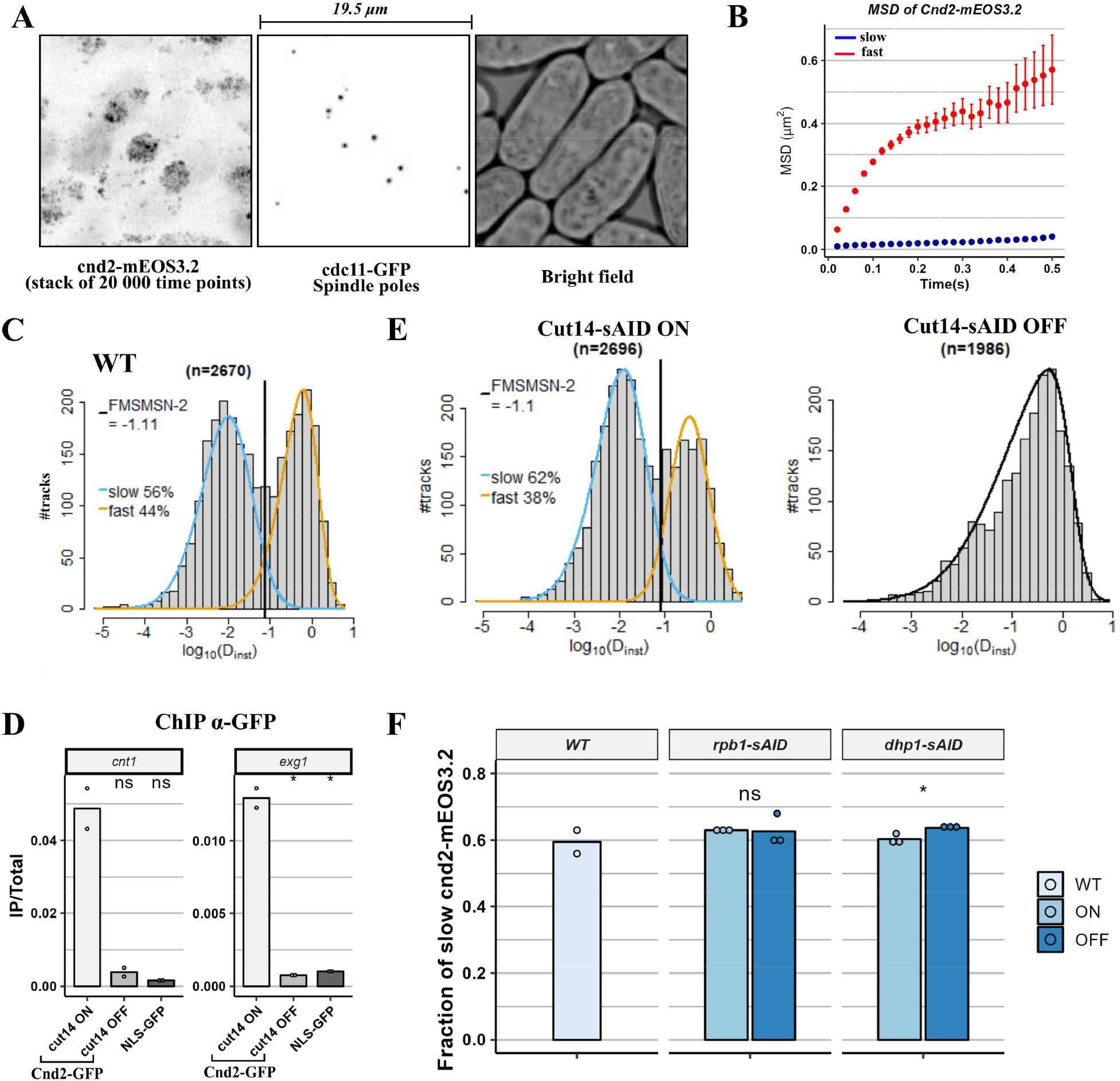
RNAP2 plays no major role in the steady-state level of condensin on chromatin. (A) Representative images of SPT-PALM acquisition on metaphase arrested fission yeast cells expressing Cnd2-mEOS3.2. Left, stack of Cnd2-mEOS3.2 during a 20 000-time series (27.7 Hz) using a TIRF illumination. Middle, position of the SPBs (spindle poles) labelled by Cdc11-GFP. Right, brightfield acquisition. (B) Median (+/- SE) mean-square displacement of the two populations of Cnd2-mEOS3.2 identified during metaphase as slow (blue) or fast (red) in WT cells (2670 tracks). (C) Distribution of the apparent diffusion coefficient (µm^2^.s^-1^) of 2670 tracks from metaphase arrested WT cells identified from one representative SPT-PALM acquisition. The vertical line represents the optimal log_10_(D_inst_) separating the two populations of molecules according to a skewed-Gaussian mixture model (FMSMSN). The fitted gaussians are represented in blue and orange for slow and fast populations respectively. The fraction of molecules belonging to each population is indicated on the left of the panel. (D) ChIP α-Cnd2-GFP after 2 hours of NaOH (ON) or IAA (OFF) treatment on either *cut14-sAID cnd2-GFP* or *NLS-GFP* cells arrested in metaphase. qPCRs were performed on two biologically independent experiments on centromere I (*cnt1)* or the condensin-enriched *exg1* gene. For statistical analysis t.test was used to compare the OFF and ON conditions: * p.value < 0.05. (E) Same as in (C) but for metaphase arrested Cut14 ON and OFF cells. The same model fitting was applied to both conditions but a single population of molecules was identified in the Cut14-OFF condition. Representative results of at least 2 independent experiments. (F) The fraction of slow Cnd2-mEOS3.2 molecules identified as described in (B-C) was calculated for WT, *rpb1-sAID* and *dhp1-sAID* metaphase arrested cells. Each point corresponds to a biologically independent experiment and at least two consecutive acquisitions (1000- 4000 tracks). t.test was used to compare the OFF and ON conditions: * p.value < 0.05.

### Active RNAP2 hinders the folding of chromatin into mitotic chromosomes

In fission yeast, cohesin folds chromatin into TADs of 30-50 kb during interphase^49,50^, while condensin shapes larger intrachromosomal domains in mitosis (median size ∼ 300-500 kb) by mediating cis contacts over longer distances ranging from ∼ 70 kb to ∼ 1 Mb^8,38,50^. To determine the consequences of transcribing RNAP2 on condensin activity, we assessed the impact of Rpb1 and Dhp1 on chromatin folding in mitosis, using chromosome conformation capture (Hi-C). We found that depleting Rpb1 from metaphase-arrests increased the frequencies of intrachromosomal contacts between 20 kb and 2 Mb (Fig. 3A, left panel), suggesting enhanced cohesin and/or condensin activity. The average size of chromatin loops can be inferred by studying the slope of the distance law, the maximum value closely matching the average length of loops^51^. Using this metric, we further found that the average size of loops in metaphase was increased upon depletion of Rpb1 (Fig. 3A, right panel). In contrast, depleting Dhp1 decreased the frequencies of intrachromosomal contacts between 100 kb and 1 Mb and reduced the average size of chromatin loops (Fig. 3B). Thus, Rpb1 limits chromatin folding in metaphase while Dhp1 promotes the formation of longer loops in the range of condensin activity. In line with this, comparing Hi-C maps between the ON and OFF conditions revealed that removing Rpb1 erased large TADs (Fig. 3C and S3A) to the benefit of long-range interactions across their borders where both Rpb1 and Cnd2-GFP were strongly reduced (Fig. 3D and S3A). Small TADs appeared reduced as well (Fig. 3C). In contrast, degrading Dhp1 decreased the frequencies of intrachromosomal contacts between large TADs (Fig. 3E and S3B) while increasing Cnd2-GFP and Rpb1 levels at the cognate borders (Fig. 3F). Similar results obtained from a second set of biological replicates are shown in Figure S3C-F. Thus, Rpb1 hinders the folding of chromatin in metaphase while Dhp1, in contrast, promotes condensin-mediated long-range 3D contacts. Since depleting fission yeast condensin is sufficient to erase large TADs characteristic of metaphase chromosomes^8^, the concurrence between a gain of contacts across a TAD boundary and a local reduction of Cnd2-GFP occupancy (Rpb1-OFF), and vice-versa (Dhp1-OFF), might therefore reflect condensin’s accumulation at a RNAP2- dependent barrier to loop formation.

**Figure 3.**
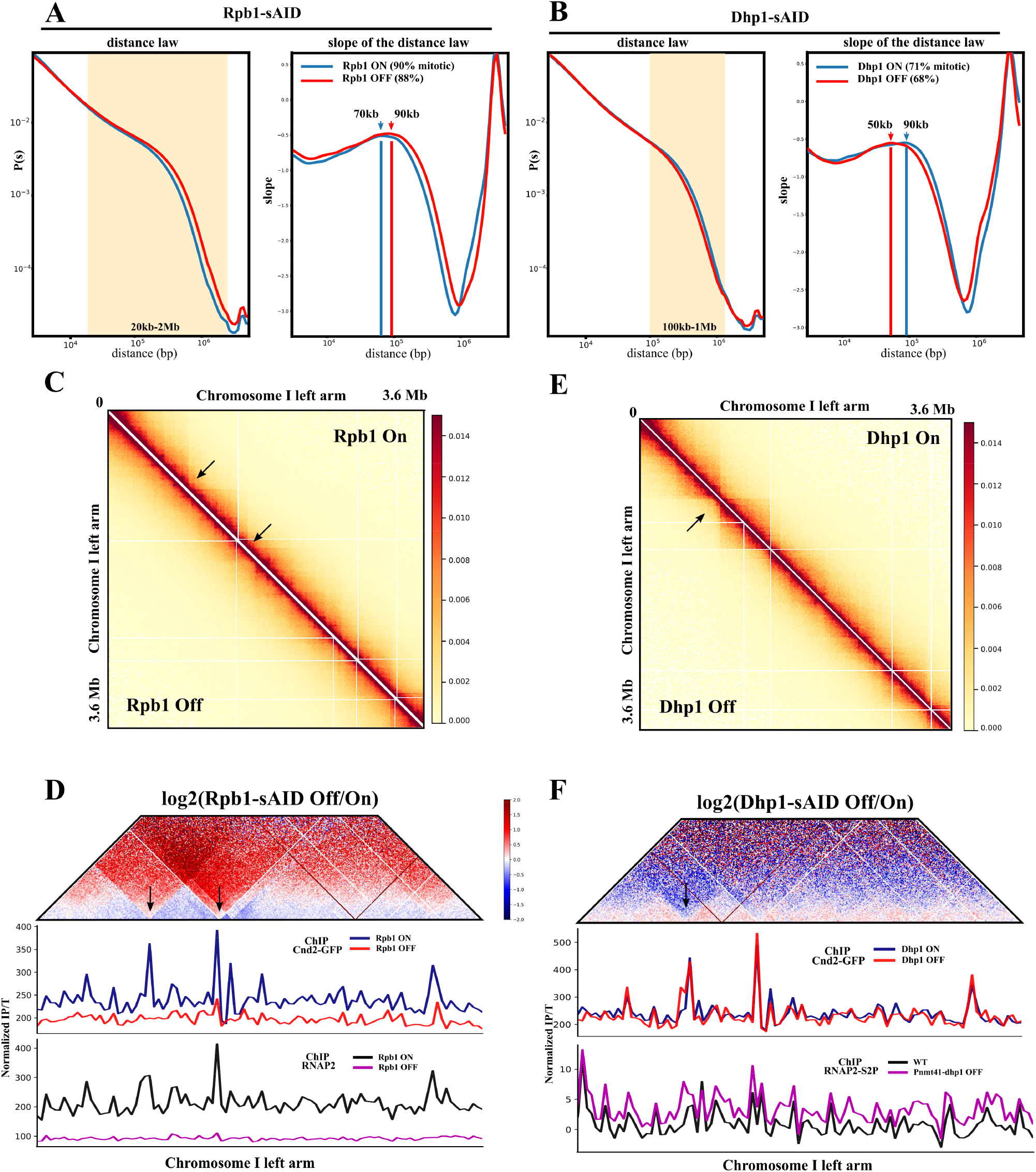
Active RNAP2 hinders chromatin folding by condensin in metaphase. (A-B) Hi-C contact probability curve P(s) as a function of distance and its corresponding slope in Rpb1-ON and Rpb1-OFF conditions (A) or in Dhp1-ON and Dhp1-OFF conditions (B). Arrows indicate the estimated size of the maximum in the slope. (C) Hi-C contact maps of Rpb1-ON and Rpb1-OFF at 10kb resolution on the left arm of chromosome I. Black arrows point to domain borders that disappear in Rpb1-OFF. (D) log2 differential map of Rpb1-OFF/Rpb1-ON at 10kb resolution on the left arm of chromosome I (Top) and tracks of the cal-ChIPseq of Cnd2-GFP and RNAP2 presented in Fig. 1 on the same region. Black arrows are the same as in (C) (E-F) Same as in (C) and (D) but for Dhp1-OFF/ Dhp1-ON. Black arrow indicates a border that is strengthened in the Dhp1-OFF condition.

### Active RNAP2 creates barriers for chromatin loop formation by condensin

We assessed whether condensin local enrichment correlated with boundaries. Using the insulation score and MACS2, we scanned the genome for any boundary and high-occupancy condensin peak, respectively. We identified 280 boundaries and 290 condensin peaks in RPB1- ON metaphases, 56 of which overlapped (Fig. 4A, red dots). This limited overlap is consistent with previous studies^8,38^ and can conceivably be explained, at least in part, by the presence within the data sets of cohesin boundaries, since small TADs formed by the latter persist in mitosis in fission yeast^50^. The insulating strength of a border correlated positively with condensin enrichment, notably when associated with a condensin peak (Fig. 4A). Aggregating Hi-C signals around two adjacent high-occupancy condensin sites confirmed that they define intrachromosomal domains (Fig. 4B). Depleting Rpb1 increased the frequencies of intrachromosomal 3D contacts across those boundaries while the insulation of condensin domains was strengthened in Dhp1-OFF (Fig. 4B), suggesting that condensin boundaries depend on their associated levels of active RNAP2. We next assessed how the changes in insulation compared with condensin occupancy. Of the 280 boundaries identified in Rpb1-ON, 45% were weakened upon depletion of Rpb1, 49% remained stable and 6% appeared strengthened (Fig. 4C). Weakened boundaries showed a high condensin peak in Rpb1-ON which was virtually erased in Rpb1-OFF (Fig. 4D), while stable and strengthened boundaries respectively exhibited unchanged and slightly increased condensin occupancy relative to the surrounding basal signal (Fig. 4D). Conversely, boundaries gaining in insulation (19%) in Dhp1- OFF gained in condensin occupancy, but stable (52%) or weakened (29%) borders exhibited no clear change (Fig. 4C-D). Thus, active RNAP2 creates insulating barriers at highly expressed genes where the accumulation of condensin matches the insulation strength.

**Figure 4.**
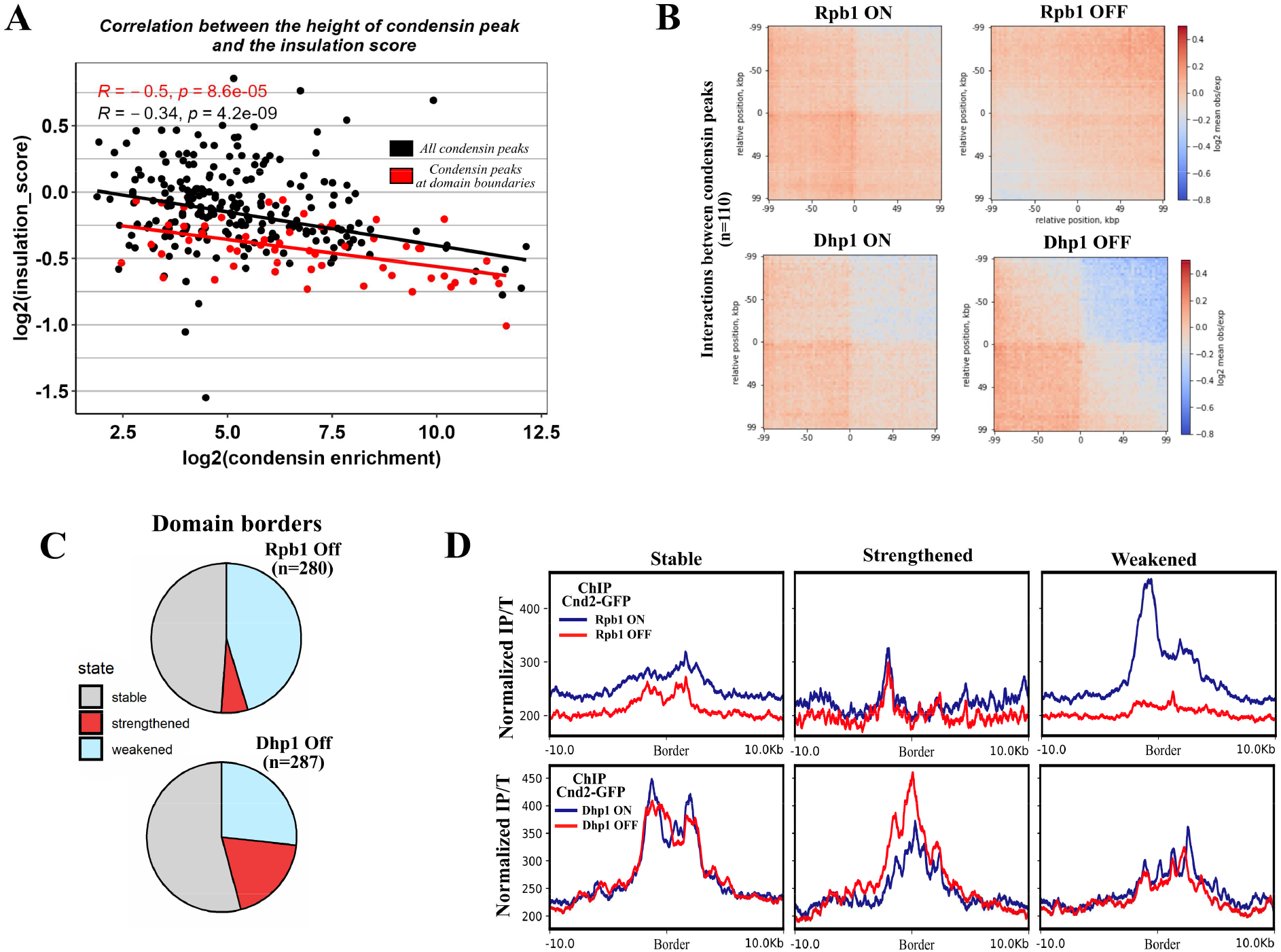
Active RNAP2 creates barriers for chromatin loop formation by condensin. (A) Distribution of the insulation score determined from Hi-C maps as a function of condensin enrichment at peaks identified by MACS2. Condensin peaks were considered overlapping with a border when they were located within the 3kb bin defining the border and are shown in red. The Pearson correlation coefficient r and its associated p.value are plotted in black for all peaks, in red for condensin peaks associated with borders. The linear regression for each population is drawn. (B) Off-diagonal pileup of Hi-C signal at n=110 condensin sites identified in the Rpb1-ON condition separated by a distance range of 100-300kb in Rpb1-ON or Rpb1-OFF and Dhp1-ON or Dhp1-OFF. Signals are shown as a function of the observed/expected log2 ratio. (C) Domain boundaries were identified with the insulation score and compared between the OFF and ON samples. A border was considered stable if the log2 ratio OFF/ON was between −0.58 and 0.58, and strengthened or weakened otherwise. (D) Metagene profiles of the mean normalized Cnd2-GFP cal-ChIPseq signal at the border classes shown in (C) in the indicated conditions.

### Transcription termination limits the strength of RNAP2 barriers to condensin

To understand how Dhp1 controls border strength, we sorted protein coding genes according to the change in condensin binding upon Dhp1 depletion using K-means clustering (Fig. 5A left panel) and analysed those genes for specific features. Among five clusters, cluster 1 revealed the strongest increase in condensin level within gene bodies (Fig. 5A, left panel). Strikingly, in Dhp1-ON, these cluster 1 genes exhibited high levels of condensin at their 3’ end but, unlike the other clusters, also at their 5’ ends (Fig. 5A, right panel). Cluster 1 genes were more frequently in convergent orientation (Fig. 5B) relative to the other clusters. They were also smaller and closer to their neighbouring genes (Fig. S4A-B) and displayed higher levels of active RNAP2 upstream of their 5’end (Fig. S4C). Such biases suggested that cluster 1 genes could be prone to invasion by transcriptional readthroughs.

**Figure 5.**
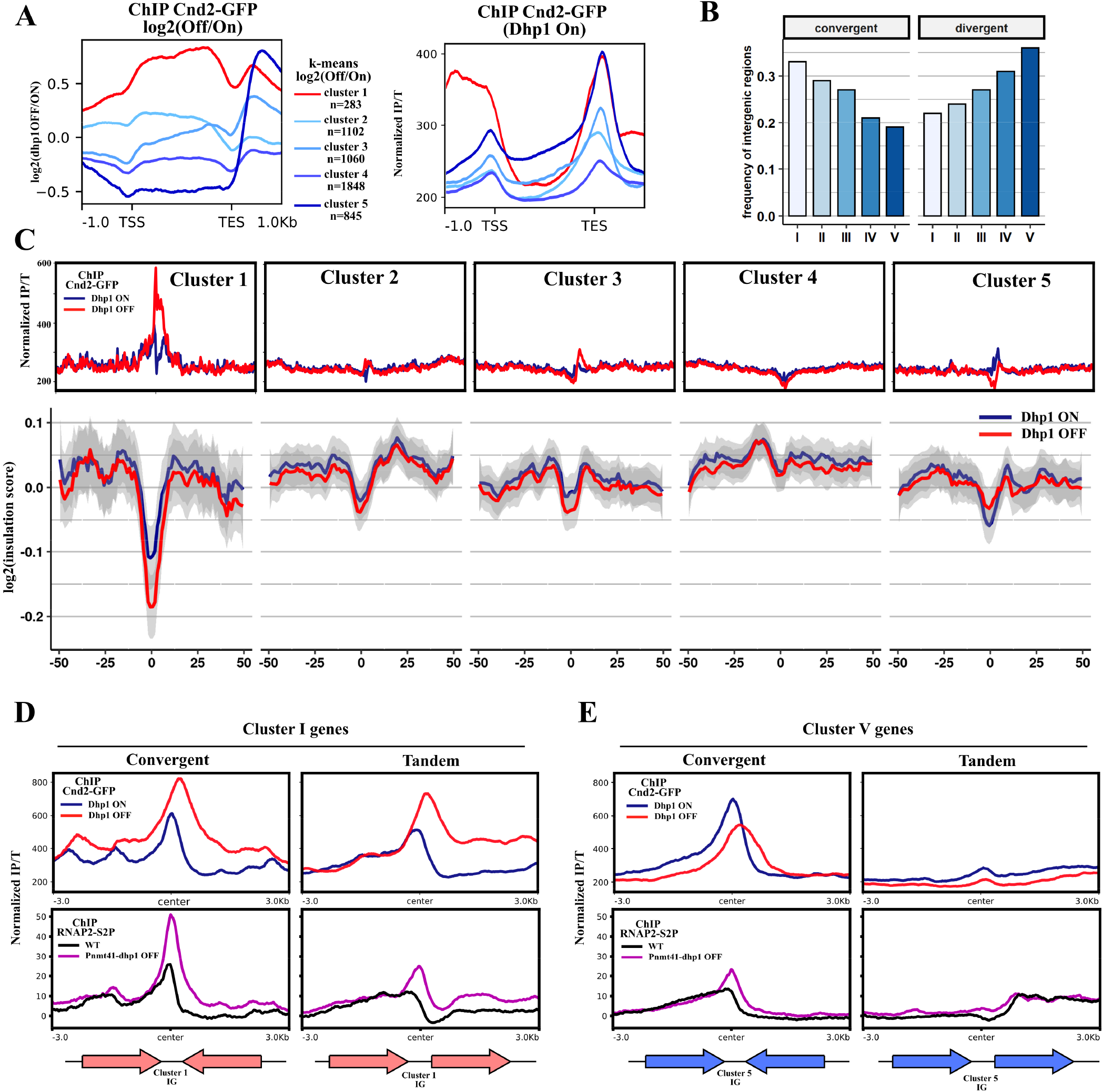
Transcription termination limits the strength of RNAP2 barriers to condensin. (A) Left panel: metagene profile of the mean log2 ratio of Dhp1-OFF/Dhp1-ON normalized Cnd2-GFP cal-ChIPseq signals, grouped after k-means clustering (k=5) at protein coding genes. Right panel: metagene profile of the mean normalized Cnd2-GFP cal-ChIPseq signal in Dhp1- ON using the same clusters. (B) Frequency of intergenic regions flanked by protein-coding genes in convergent or divergent orientations in clusters I to V defined in (A). (C) Mean normalized Cnd2-GFP cal-ChIPseq signal at clusters I to V in the indicated conditions and their corresponding mean Hi-C insulation score (bottom). The region shown is 100kb centered on the border. (D) Metagene profile of mean normalized Cnd2-GFP and RNAP2 (S2P)^44^ cal-ChIPseq-signals in Dhp1-ON or OFF samples of cluster I genes. A 6kb window centered on the intergenic region flanked by converging or tandem genes is shown. For convergent gene pairs, the most expressed gene is always on the left. (E) Same as in (D) for cluster V.

Cluster 1 genes were the most insulating in Dhp1-ON and showed the strongest gain in insulation in Dhp1-OFF (Fig. 5C), suggesting that the depletion of Dhp1 mostly strengthened pre-existing boundaries. In contrast, the other clusters showed lower insulation in both conditions. Notably, cluster 5, which displayed similar levels of condensin as cluster 1 in Dhp1- ON but only at the 3’ end (Fig. 5A), gained neither in condensin occupancy within gene bodies (Fig. 5A) nor in insulation upon depletion of Dhp1 (Fig. 5C). Considering the existing biases in cluster 1 genes which are reversed in cluster 5 (Fig. 5B and S4), we assessed whether transcriptional readthroughs could explain their different sensitivities to Dhp1. To this end, we selected the nearest facing neighbour of each gene of cluster 1 and assessed the condensin and RNAP2 ChIP-seq profiles of every gene pairs (Fig. 5D). This clearly showed that depleting Dhp1 caused transcriptional readthroughs across their common intergenic region and increased RNAP2 occupancy in the body of the less transcribed downstream gene (Fig. 5D, lower panels), a likely consequence of the invasion of the latter by reading-through polymerases. The condensin peak initially proximal to the TES of the upstream gene became higher and larger upon depletion of Dhp1 and was displaced in the direction of the strongest transcription, encroaching on the body of the downstream gene (Fig. 5D, upper panels). The same analysis applied to cluster 5 revealed readthroughs of lower amplitude and no visible invasion of condensin into the downstream gene (Fig. 5E). Thus, these data suggest that Dhp1 dampens the strength of condensin borders by ensuring an efficient termination and removal of RNAP2 from chromatin.

### RNAP2 hinders condensin-dependent mitotic chromosome segregation in anaphase

The formation of chromatin bridges in anaphase caused by condensin deficiency conceivably stems from the fact that condensin orientates the activity of Topoisomerase II towards the decatenation of chromosomes and sister-chromatids^52,53^ and confers to chromosome arms the elasticity to withstand the spindle traction forces^54^. If condensin achieved these tasks by folding chromatin, depleting Rpb1 or Dhp1 was expected to respectively lower or increase the frequency of chromatin bridges caused by a mutation in condensin. We and others previously observed a rescue of accurate chromosome segregation in condensin mutant cells when a component of the transcriptional co-activator Mediator was impaired^55^ or upon pharmacological inhibition of transcription^25^. We therefore measured the effect of depleting Rpb1 or Dhp1 on the frequency of chromatin bridges caused by the thermosensitive *cut3-477* mutation in the Cut3^SMC4^ subunit of condensin^2^, which reduces condensin binding at the restrictive temperature of 36°C^33^. Consistent with previous reports^2,55^, ∼80% of *cut3-477* cells in anaphase displayed a chromatin bridge at 36°C (Fig. 6A-B). The frequency sharply dropped to ∼ 40% when Rpb1 was depleted in *cut3-477* mutant cells (Fig. 6B). Depleting Dhp1, in contrast, almost doubled the frequency of chromatin bridges caused by the *cut3-477* condensin mutation at the semi-permissive temperature of 32°C (Fig. 6C), as expected. To validate these observations, we next assessed the impact of the DNA-binding transcription factor Fkh2, which together with Ace2 drives the transcription of most of the mitotically expressed protein-coding genes where condensin accumulates in mitosis^38,56^. Consistent with the rescue effect by Rpb1 depletion, deleting *fkh2* restored the growth of *cut3-477* mutant cells at 36°C (Fig. 6D). Thus, these data extend the idea that Rpb1 and Dhp1 respectively antagonises and facilitates the functioning of condensin from metaphase to anaphase by affecting its ability to fold chromatin and by extension to support accurate chromosome segregation.

**Figure 6.**
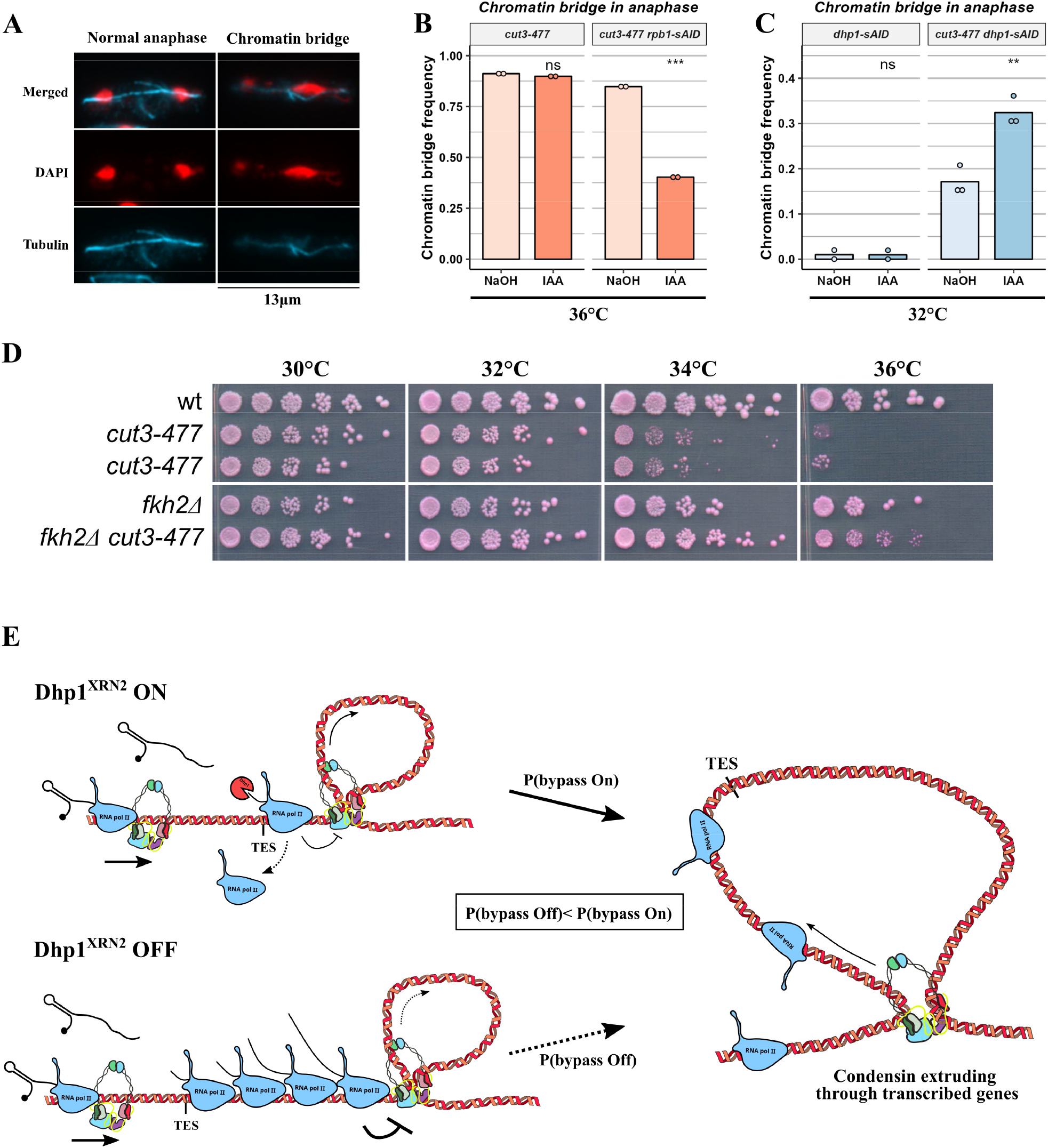
RNAP2 hinders condensin-dependent mitotic chromosome segregation in anaphase. (A) Representative images of normal and defective anaphases in fission yeast cells. (B-C) Indicated cells were fixed and processed for immunofluorescence against α-tubulin. Anaphase cells showing a mitotic spindle > 5 µm in length were selected and chromosome segregation assessed. Shown are frequencies calculated from 2 or 3 replicates with n = 100 anaphases per experiment. (D) Serial dilution spot assays (5-fold) of the indicated *S. pombe* strains. Images are taken after 60 hours of growth at the indicated temperatures. (E) Model of condensin activity at highly transcribed genes. In wild-type, loop-extruding condensin accumulates at chromosomal sites densely bound by RNAP2, such as the 3’ end of genes near the TES. Condensin can eventually bypass this domain-defining barrier with an indeterminate probability. When Dhp1^XRN2^ is depleted from metaphase cells, RNAP2 transcribes through the TES, pushing condensin complexes further downstream. At gene dense regions, Dhp1^XRN2^ depletion causes reading through RNAP2 to invade downstream genes, creating larger and less permeable barriers that further stall condensin and lowers its ability to extrude beyond this region.

## Discussion

Here we investigated the enrichment of fission yeast condensin at genes highly transcribed by RNAP2, and the functional consequences of such an accumulation on chromatin folding in metaphase and accurate chromosome segregation in anaphase. By rapidly and acutely depleting Rpb1 and Dhp1, we modulated the occupancy and the location of transcribing RNAP2 in metaphase and collected compelling experimental evidence showing that RNAP2 does not control the steady state level of condensin binding but instead creates transcriptional barriers that accumulate condensin and hinder condensin-mediated chromatin folding and chromosome segregation. Our results are in perfect agreement with the stalling of translocating bacterial SMCs against transcriptional barriers^57,58^, and fully consistent with the way transcription interferes with cohesin localisation and loop-formation activity in various systems^17–19,59,60^. Thus, given the large body of studies supporting loop extrusion by cohesin in vivo^9^, and by all eukaryotic SMCs in vitro^10–14^, we argue that transcription is a barrier for chromatin folding by condensin in vivo as indirect evidence that condensin folds mitotic chromosomes by loop extrusion (Fig. 6E).

Our finding that the steady state level of chromosomal condensin is not modified by the acute depletion of Rpb1 sheds a new light on the possible molecular determinants of condensin loading sites within chromatin. First, it demonstrates that Rpb1, the core component of RNAP2, is not required for condensin loading. Second, it challenges the idea that unwound or positively supercoiled DNA structures generated by transcription could recruit condensin ^39,61^ as they are unlikely to survive the depletion of Rpb1^62^. On the other hand, DNA-bound transcription factors and nucleosome-depleted promoters can persist in mitosis despite transcriptional repression^63,64^, raising the possibility that they may underly the Rpb1- independent localisation of condensin in the vicinity of some gene promoters. Further study is needed to assess whether or not it could be the case.

Regarding the mechanism by which RNAP2 impinges upon condensin, the most parsimonious hypothesis should explain how RNAP2 positions condensin along the genome while antagonising its chromatin folding activity. The moving barrier model^19,58^ fulfils these criteria. Indeed, the stalling of loop extruding condensin molecules upon head-on collision with a transcriptional barrier formed in a Rpb1-dependent manner (Fig. 6E) provides a straightforward explanation to the redistribution of condensin, the gain of long-range intrachromosomal contacts in metaphase and the weakening of condensin boundaries in Rpb1-depleted metaphases. The removal of a transcriptional hindrance to condensin-mediated chromatin folding is also consistent with the rescue of accurate chromosome segregation observed in condensin mutant cells upon depletion of Rpb1. Such a mechanistic model also convincingly explains our experimental observations in Dhp1-depleted cells (Fig. 6 6E, as expected from the control exerted by Dhp1 over Rpb1^44^. The loss of Dhp1 leads to (1) an enlargement of the domains occupied by active RNAP2, (2) a rise in condensin occupancy in cis that positively correlates with boundary strength, particularly at a subset of convergent genes separated by short intergenic regions, (3) reduced frequencies of intrachromosomal contacts and (4) an increased hindrance on condensin-mediated chromosome segregation. A similar mechanism likely explained the accumulation of condensin at a subset of tRNA genes that we previously observed when the RNAP3 transcription termination factor Sen1 was deleted^21^. On the other hand, the possibility that Dhp1 and/or Rpb1 could impact condensin-mediated diffusion capture appears less likely. Since high-occupancy condensin binding sites are postulated to mediate diffusion capture^22,23^, it is difficult to imagine how the mere redistribution of condensin upon depletion of Rpb1 could benefit the accurate segregation of chromosomes in anaphase in a condensin mutant background, and reciprocally depleting Dhp1 be detrimental to the folding of chromatin by diffusion capture while it increases condensin occupancy at those sites.

It is therefore tempting to speculate that the broad and basal condensin-ChIP-seq signal observed in a wide range of species correspond to translocating condensin complexes caught in the process of loop-extrusion. The slight increase in condensin occupancy in Dhp1-depleted metaphases might stem from an increased residence time of such active condensin complexes trapped in between converging reading-though polymerases, among other possible mechanisms. Along this line, it is unclear why Dhp1-depletion had no apparent impact on the frequencies of contacts in the range size of cohesin in our Hi-C data. This might be due to the fact that our experiments were performed in metaphase and/or to varying sensitivities to Dhp1 loss since condensin and cohesin respond differently to single obstacles in loop-extrusion assays^65^.

Although loop-extruding condensin has been observed traversing DNA-bound isolated obstacles of sizes ranging from tens to 200 nm in in-vitro assays^65^, it has also been reported that engineered arrays of tightly bound proteins impair in *cis* the activity of condensin in budding yeast^20^. Thus, features associated with tracks of RNAP2, be it proteins, nascent RNA, chromatin modifications, specific DNA structures and/or changes in the local chromatin fiber rigidity are as many possible candidates to stall condensin. Whatever the molecular determinants, the fact that read-through transcripts are produced in wild-type fission yeast cells^66,67^ and that condensin accumulation and insulation are found more often at closely spaced convergent genes points towards efficient transcription termination as a key player in the folding of transcribed chromatin by condensin complexes. In that context, the downregulation of transcription taking place upon mitotic entry in metazoans might provide an increase in fitness for the folding of large genomes by loop extruding condensin.

## Materials and Methods

Media, growth, maintenance of strains and genetic methods were as described^68^. Standard genetics and PCR-based gene targeting method were used to construct *S. pombe* strains and sanger sequencing was performed to confirm the insertions. Dhp1 and Rpb1 were tagged at their C-terminus with 3x sAID repeats and the degron alleles transferred into *osTIR1-F74A*^69^ expressing genetic backgrounds by crossing. The *cnd2-GFP* and degron alleles are expressed under the control of their natural promoter at their native chromosomal location. Strains used in this study are listed in Table S1. For metaphase arrests, cells expressing the APC/C co- activator Slp1 under the control of thiamine-repressible promoter *nmt41* were cultured in synthetic PMG medium at 32°C and arrested in metaphase for 3 hours at 32°C by exposure to 20 µM thiamine. Mitotic indexes were determined by scoring the percentage of cells exhibiting Cnd2-GFP fluorescence in their nucleoplasm^3^. Liquid cultures of cells expressing either Dhp1-sAID or Rpb1-sAID treated with thiamine to induce their metaphase arrest were exposed to 100 nM 5-adamantyl-IAA (or NaOH as control) for 1h or 30 minutes (min), respectively, before the end of the 3h metaphase arrest.

### Chromosome segregation assay

Cells were grown to exponential phase in PMG liquid medium at 32°C (for dhp1-sAID) or grown in YES+A liquid medium at 30°C and shifted for 2.5 h at 36°C (for rpb1-sAID). Before collecting cells, liquid cultures were exposed to 100 nM 5-adamantyl-IAA or NaOH as control (1h before collecting for dhp1-sAID or 30 min for rpb1-sAID). 2.10^7^ cells were fixed in cold methanol and stored at −20°C. Cells were processed for immunofluorescence by washing three times with PEM (100 mM PIPES, 1mM EGTA, 1mM MgSO4, pH 6.9) with the last wash performed on a wheel at room temperature for 30 mn to rehydrate cells. Cells were digested with 0.4mg of zymoliase 100T (nacalai tesque) in PEMS (PEM + 1.2M Sorbitol) for 30 min at 37°C in a waterbath. Cells were washed twice with cold PEMS, and incubated for ∼1-2 min in PEMS + 2% Triton at room-temperature. Cells were pelleted, washed with PEM and resuspended in PEMBAL (PEM, 1% BSA, 100mM Lysine-HCl, 0,1% sodium azide) on a wheel at room temperature for 30 min. Cells were resuspended in 100 µl PEMBAL with 1/50 TAT1 antibody and incubated on a wheel overnight at 4°C in the dark. Cells were washed three times with PEMBAL, incubated in 100 µl PEMBAL with 1/400 anti-mouse AlexaFluor488 fluorescent antibody for 2h on a wheel at RT. Cells were washed a final time in PEMBAL and resuspended in PEM+0,5 µg/ml DAPI. 4 µl were placed on a slide, covered by a coverslip and observed with Zeiss Axioscope A.1, objective Apopchromat 63x 1.4NA. Chromatin bridges were scored manually.

### Calibrated-ChIP and sequencing

Calibrated ChIP was performed as described previously^33^. Briefly, fission yeast cells expressing either Cnd2-GFP or NLS-PK9-GFP and arrested in metaphase by the depletion of Slp1 were fixed with 1% formaldehyde for 25 min, quenched with glycine 0.125 M final, washed twice with PBS, frozen in liquid nitrogen and stored at −80°C until use. *Saccharomyces cerevisiae* cells expressing Smc3-GFP, used for internal calibration, were grown in Yeast Peptone Dextrose liquid medium at 30°C in log phase and fixed with 2.5% formaldehyde for 25 min. To perform calibrated ChIPseq against Cnd2-GFP or RNAP2, fixed fission yeast and budding yeast cells were mixed at a ratio of 5:1 prior to lysis with Precellys® (Bertin). Anti-GFP and anti-RNAP2 ChIPs were performed using the A111-22 and 8WG16 antibodies, respectively. Libraries were prepared using NEBNext® Ultra™ II DNA Library Prep Kit for Illumina® according to the manufacturer’s instructions. DNA libraries were size-selected using Ampure XP Agencourt beads (A63881) and sequenced paired-end 150 bp with Novaseq S6000 (Novogene®). To make ChIP-seq libraries for Rpb1-sAID depletion, IP and Total fractions from three independent biological experiments were pooled.

### Hi-C

Fission yeast cells, expressing Cnd2-GFP and arrested in metaphase by the depletion of Slp1 were fixed with 3% formaldehyde for 5 min at 32°C followed by 20 min at 19°C, washed twice with PBS, frozen in liquid nitrogen and stored at −80°C. 2.10^8^ cells were lysed in ChIP lysis buffer with Precellys® (Bertin). Lysates were centrifuged 5000 g at 4°C for 5 min and pellets were resuspended once in 1 mL lysis buffer and twice in NEB^®^ 3.1 buffer. SDS was added to reach 0.1% final and samples were incubated for 10 min at 65°C. SDS was quenched on ice with 1% Triton X-100 and DNA digested overnight at 37°C with 200 Units of *Dpn*II restriction enzyme. Samples were incubated at 65°C for 20 min to inactivated *Dpn*II. Restricted-DNA fragments were filled-in with 15 nmol each of biotin-14-dATP (cat. 19524016, Thermofisher), dTTP, dCTP and dGTP, and 50 units of DNA Klenow I (cat. M0210M, NEB) for 45 min at 37°C. Samples were diluted in 8 ml of T4 DNA ligase buffer 1X and incubated 8 hours at 16°C with 8000 Units of T4 DNA ligase (NEB). Crosslinks were reversed overnight at 60°C in the presence of proteinase K (0.125 mg / ml final) and SDS 1% final. 1 mg of proteinase K was added again and tubes were further incubated for 2 hours at 60°C. DNA was recovered by phenol-chloroform-isoamyl- alcohol extraction, resuspended in 100 µl TLE (Tris/HCl 10 mM, 0.1 mM EDTA, pH8) and treated with RNAse A (0.1 mg / ml) for 30 min at 37°C. Biotin was removed from unligated ends with 3 nmol dATP, dGTP and 36 Units of T4 DNA polymerase (NEB) for 4 hours at 20°C. Samples were incubated at 75°C for 20 min, washed using Amicon® 30k centrifugal filters and sonicated in 130 µl H_2_O using Covaris® S220 (4 min 20°C, duty factor 10%, 175W peak power, 200 burst per cycle). DNA was end-repaired with 37.5 nmol dNTP, 16.2 Units of T4 DNA polymerase, 54 Units of T4 polynucleotide kinase, 5.5 Units of DNA Pol I Klenow fragment for 30 min at 20°C and then incubated for 20 min at 75°C. Ligated junctions were pulled-down with Dynabeads® MyOne™ Streptavidin C1 beads for 15 min at RT and DNA ends were A-tailed with 15 Units of Klenow exo- (cat. M0212L, NEB). Barcoded PerkinElmer adapters (cat. NOVA- 514102) were ligated on fragments for 2 hours at 22°C. Libraries were amplified with NextFlex PCR mix (cat. NOVA-5140-08) for 5 cycles, and cleaned up with Ampure XP Agencourt beads (A63881). Hi-C libraries were paired-end sequenced 150bp on Novaseq6000 (Novogene®).

### SPT-PALM

Cells were cultured at 30°C in filter-sterilized PMG for 48h and shifted to 25°C, in exponential phase, for 5 hours. 3.5 h before the beginning of the acquisition, thiamine 20 μM was added to the culture to repress transcription of *nmt41-slp1* gene and arrest cells in metaphase. 10 min before the acquisition, 2 mL of culture was taken, centrifuged 15 sec at 10 000 g at room temperature and resuspended to reach 5.10^8^cells/mL. Cells were transferred to an agarose pad as described below. Coverslips (Marienfeld 0107052 22x22mm No. 1.5H) were washed 10 min in acetone, two times 5 min in ethanol, 10 min in KOH 1M, five times in Milli Q water and finally dried with a flame. Coverslips showing a defect or a carbon deposit after the drying step were discarded. On a clean glass slide (RS BPB018, frosted end), two pieces of double-sided tape were placed to fix the CoverWell imaging chamber (Grace Bio-labs 635021, 25x25mm). 1 mL of PMG 2% Agarose (invitrogen 16500) supplemented with 20 μM thiamine was poured in the CoverWells. 5 μL of fresh cell suspension (5.10^8^ cells/mL) were deposited on the pad and covered with a KOH-treated coverslip, sealed with nail varnish and immediately used for acquisition. We depleted Rpb1-sAID for 30 min, Dhp1-sAID for 1h and Cut14-sAID for 4h prior to acquisition. SPT-PALM microscopy was performed with a Zeiss Elyra 7 system, equipped with a TIRF setup and controlled by Zen Black software. An oil immersion 100x objective α Plan-Apochromat 1.46NA, and an EMCCD (iXon EMCCD 897 Ultra, pixel size=16μM) was used for the acquisition. Cnd2-mEOS3.2 acquisition was done in a field with 5-20 living yeast cells, using a TIRF angle around 66° which should correspond to a penetration of 100-150 nm. A 405 nm laser was used for photoconversion of mEOS3.2 with constant activation and low intensity (0.015%, 0.4-1.3 µW as measured by Optical Power Meter 100D, slide S170C with an epifluorescence illumination). Excitation of converted mEOS3.2 by the 561 nm laser was done with an exposure time of 20 ms and 20% power (5.6 mW in epifluorescence). Time Series of 20 000 cycles without interval and with Definite Focus 2 (Zeiss, continuous) were performed. During the acquisition the 405 nm laser power was slightly adjusted to always see enough particles. With these settings, the framerate was 33.6 ms. Once the time series was finished, GFP (Cdc11-GFP, SPB/centrosome) and brightfield acquisitions were done in a z-stack.

### SPT-PALM data analysis

Cells showing two separated Cdc11-GFP dots were included in the Region Of Interest (ROI). Trackmate^70^ was used on Fiji^71^ for segmentation with the LoG detector set at diameter = 1 μm and quality threshold at 100. Track reconstruction was done by the Simple LAP tracker with a Linking max distance and a gap closing distance of 0.8 μm and a gap-closing max frame of 3. Path between 5 and 50 spots were exported and analysed with both spot-ON^48^ and sptPALM viewer^46^. The spot-ON parameters were as described in ^45^ and for sptPALM viewer, on matlab we ran the script, imported the csv and set the maximal distance to 0.8 μm and minimal frame number to 5. Trajectory were further analysed in R as described^46^.

### Hi-C data analysis

Alignments were performed using *hicstuff* version 3.1.2 with options (-d –D –m iterative –e DpnII) after merging.fq.gz files of the same sample from different sequencing runs. Matrices were visualized and compared with *hicstuff view* after subsampling to the same number of contacts. For log2 differential maps, all three chromosomes were represented at 20kb binning. To visualize individual matrices, chrI :100000-3700000 was viewed at 10kb binning. Insulation scores (IS) at protein coding genes were determined using cooltools^72^ with a binning of 3kb and a window of 9kb and plotted with R. Border were identified in cooltools in all the conditions with the IS as described above. For each degron, the borders in the ON and OFF conditions were merged and the IS of their bin compared. A border was considered weakened if log2(IS-OFF/IS-ON) > 0.58, strengthened if log2(IS-OFF/IS-ON) < −0.58 and stable otherwise.

### ChIP-seq data analysis

Alignments were performed as described previously^33^. Briefly, reads were mapped using an nf-core (https://doi.org/10.1038/s41587-020-0439-x) modified pipeline with TEL2R extended ASM294v2 genome (Omnibus GEO GSE196149) for *Schizosaccharomyces pombe* mapping reads, and sacCER3 release R64-1-1 for calibrating, *Saccharomyces cerevisae* mapping reads. Deeptools2 was used to plot metagene profiles and heatmaps^73^, as well as to perform k-means clustering of bigwig data using scale-regions. Peak calling was performed on BAM files using the MACS2 software^74^. For the definition of gene-pairs in Fig. 5D, we considered as neighbours any gene oriented towards a gene of cluster I with a TES closer to 1kb from the TSS or TES of cluster I.

### Data availability

Cal-ChIP–seq and Hi-C data were deposited in the Gene Expression Omnibus under accession no. GSE236395.

## Acknowledgments

We thank T. Etheridge for the *cnd2-mEOS3.2* strain; J. Brocard for SPT-PALM analyses, F. Beckouët for critical reading of the manuscript and V. Vanoosthuyse and A. Piazza for helpful discussions. J.L. and L.C. are supported by PhD studentships from respectively the Ecole Normale Supérieure and la Ligue Contre le Cancer. This work was funded by the CNRS, ENS- Lyon and grants from the ANR (ANR-22-CE12-0035 condensinchromatin) and la Fondation ARC (PJA 20191209370) to P.B.

## Author contributions

Investigation: J.L. and L.C.; Formal Analysis: J.L., L.C. and E.C; Conceptualization, Methodology, Validation, Visualization and Writing original draft: J.L., L.C. and P.B; Supervision, Project Administration and Funding Acquisition: P.B.

## Declaration of interests

Authors declare that they have no competing interest.

## Figure Legends

**Supplementary Figure 1:**
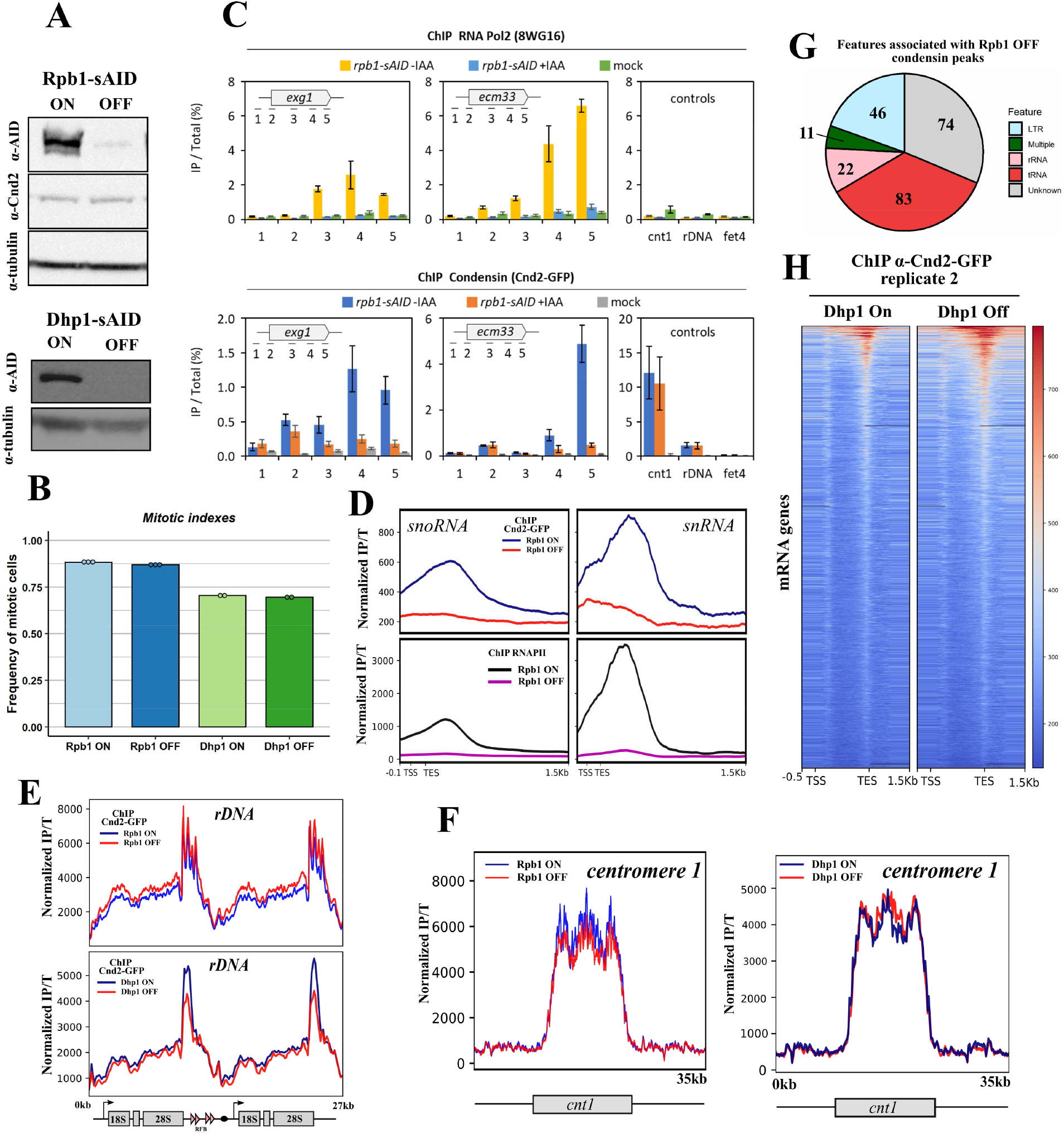
Active RNAP2 localises condensin in cis, related to Figure 1. (A) Representative western blots of total protein extracted from *rpb1-sAID* or *dhp1-sAID* metaphase arrests before and after 30 min and 1h of 5-adamantyl-IAA treatment, respectively. Tubulin is shown as a loading control. (B) Mitotic indexes, calculated as the percentage of mononucleate cells exhibiting high Cnd2- GFP fluorescent signals in their nucleus within the total cell population, used for cal-ChIP-seq in Figure 1. (C) ChIP-qPCR signal of Rpb1-ChIP (top) and Cnd2-GFP-ChIP (bottom) at two mitotically expressed protein coding genes (*exg1, ecm33*), central domain of centromere I (*cnt1*), at ribosomal DNA from the right arm of chromosome III (rDNA) and a low condensin binding site (*fet4*). Mean values from three biological replicates are shown, with standard deviation. (D) Metagene profile of the mean normalized Rpb1 and Cnd2-GFP cal-ChIPseq signal at snRNA and snoRNA genes in Rpb1-ON and Rpb1-OFF (E-F) Normalized Cnd2-GFP ChIPseq signal at the at the rDNA locus on the right arm of chromosome III (E) or at the central domain of centromere 1 (F) in Rpb1-ON and Rpb1-OFF and Dhp1-ON and Dhp1-OFF conditions. (G) Distribution of features associated with condensin peaks detected by MACS2 in Rpb1-OFF. (H) Heatmap of normalized Cnd2-GFP cal-ChIPseq signals at protein coding genes ranked by mean strength, in Dhp1-ON and Dhp1-OFF of the second biological replicate.

**Supplementary Figure S2:**
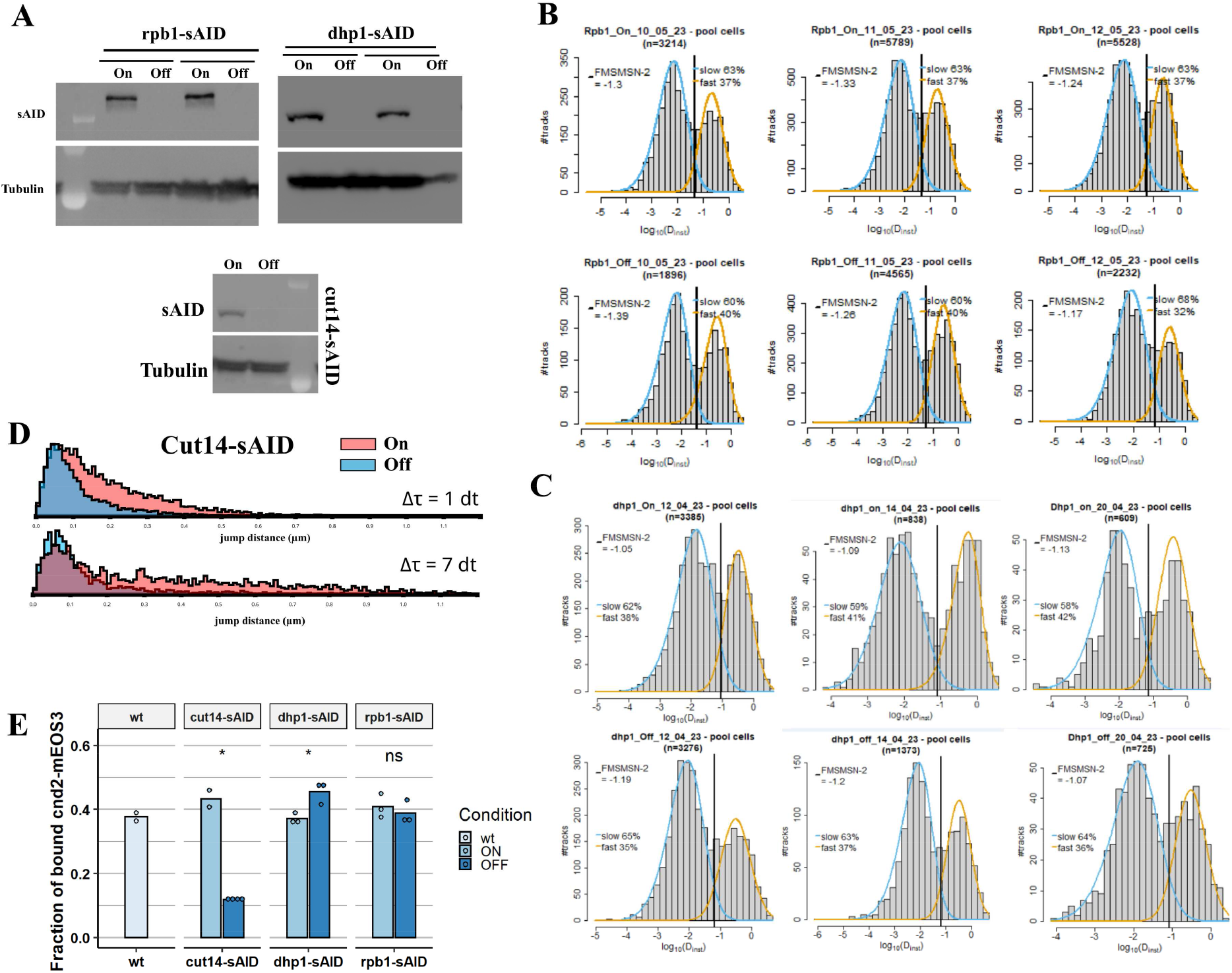
RNAP2 does not influence the steady-state level of condensin on chromatin, related to Figure 2. (A) Western blots showing the depletion of Rpb1-sAID (30 min), Dhp1-sAID (1h) and Cut14- sAID (4h) from cells used for the SPT-PALM experiments. (B) Distribution of the apparent diffusion coefficient (µm^2^.s^-1^) for the three independents *rpb1- sAID* experiments shown in Figure 2F. The vertical line represents the optimal log_10_(D_inst_) separating the two populations of molecules according to a skewed-Gaussian mixture model (FMSMSN). The fitted gaussians are represented in blue and orange for slow and fast populations respectively. The fraction of molecules belonging to each population is indicated. (C) Same as in (B) but for *dhp1-sAID* experiments. (D) Jump distance distribution calculated by spot-ON^48^ for the Cut14-sAID ON and OFF conditions. This represents the spatial distance separating two dots from the same tracks separated by Δτ ms (1 dt=33.6ms). (E) The fraction of slow Cnd2-mEOS3.2 molecules identified by spot-ON with a three-state model (Materials and Methods) quantified for WT, *cut14-sAID*, *rpb1-sAID* and *dhp1-sAID* metaphase arrested cells (same tracks as for Fig. 2F). t.test was used to compare the OFF and ON conditions: * indicates a p.value < 0.05.

**Supplementary Figure S3:**
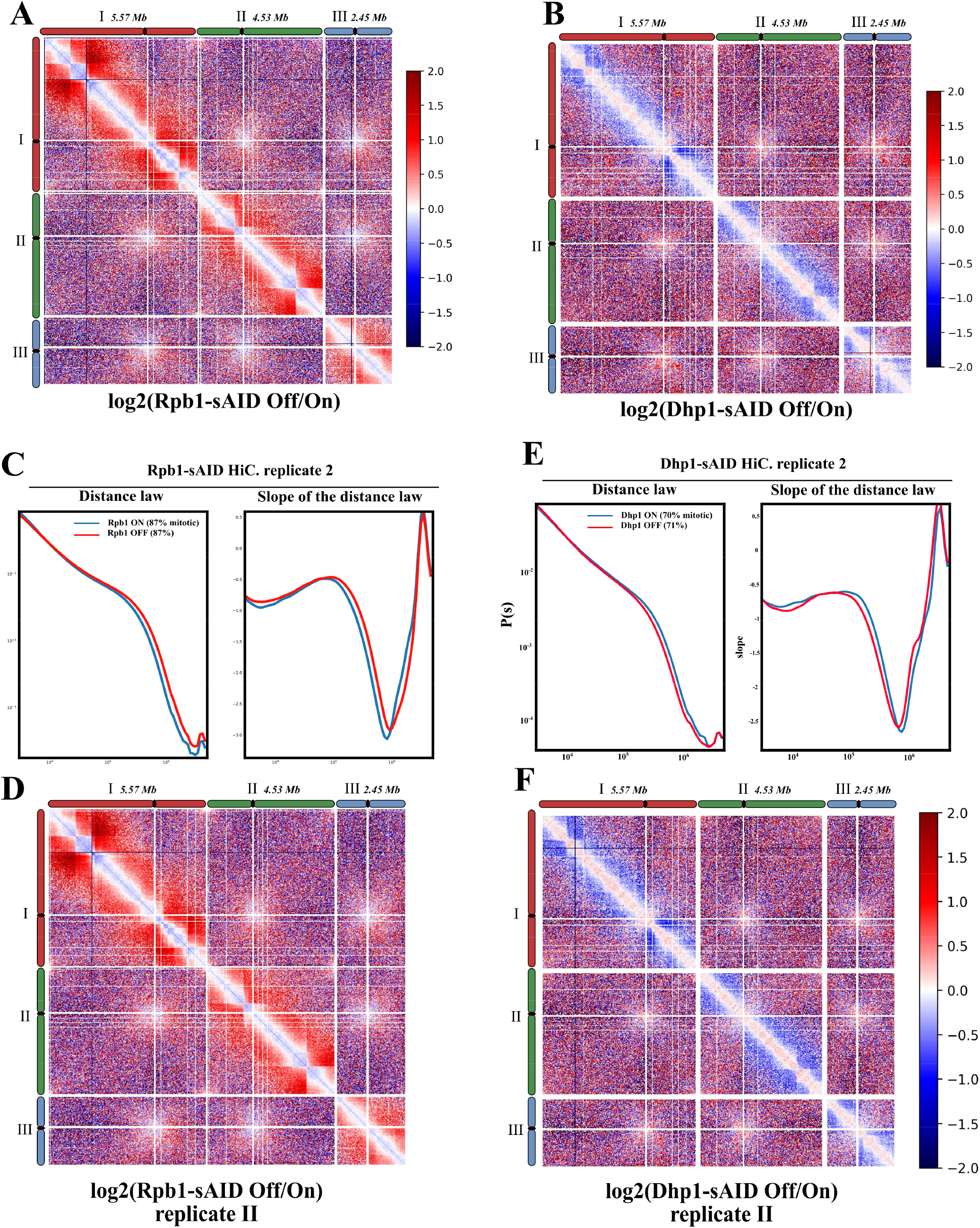
Active RNAP2 hinders chromatin folding by condensin in metaphase, relative to Figure 3. (A-B) Genome-wide log2 differential maps at 20kb resolution of (A) Rpb1-OFF/Rpb1-ON and (B) Dhp1-OFF/Dhp1-ON (C-D) Hi-C contact probability curve P(s) as a function of distance and corresponding slope and genome-wide log2 differential map at 20kb resolution (D) in Rpb1-ON and Rpb1-OFF samples of the second biological replicate (E-F) Hi-C contact probability curve P(s) as a function of distance and its corresponding slope and genome-wide log2 differential map at 20kb resolution (F) in Dhp1-ON and Dhp1-OFF samples of the second biological replicate.

**Supplementary Figure S4:**
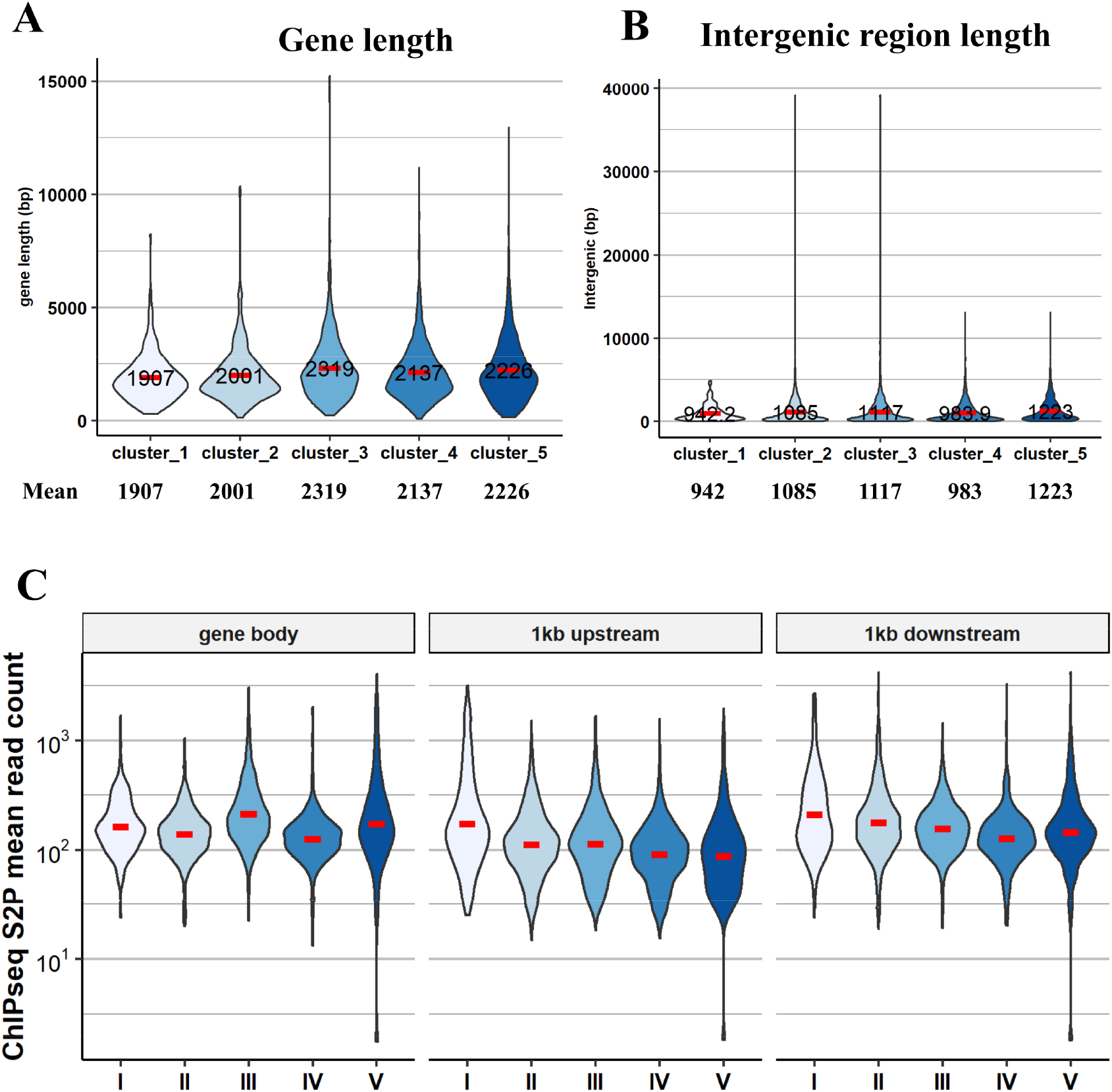
Transcription termination limits the strength of RNAP2 barriers to condensin, related to Figure 5. Violin plots of (A) gene lengths, (B) intergenic region lengths and (C) mean RNAP2 (S2P) cal- ChIPseq signals in WT in clusters defined in Figure 5A.

## Notes

### Competing Interest Statement

The authors have declared no competing interest.

